# Linguistic input drives brain network configuration during language comprehension

**DOI:** 10.1101/2020.01.22.915041

**Authors:** Ileana Quiñones, Nicola Molinaro, César Caballero-Gaudes, Simona Mancini, Juan Andrés Hernández-Cabrera, Horacio Barber, Manuel Carreiras

## Abstract

Assessing the synchrony and interplay between distributed neural regions is critical to understanding how language is processed. Here, we investigated possible neuro-functional links between form and meaning during sentence comprehension combining a classical whole-brain approach, which characterizes patterns of brain activation resulting from our experimental manipulation, and a novel multivariate network-based approach, which uses graph-theory measures to unravel the architectural configuration of the language system. Capitalizing on the Spanish gender agreement system, we experimentally manipulated formal and conceptual factors: whether the noun-adjective grammatical gender relationship was congruent or not and whether the noun gender type was semantically informative or strictly formal. Left inferior and middle frontal gyri, as well as left MTG/STG emerged as critical areas for the computation of grammatical relations. We demonstrate how the interface between formal and conceptual features depends on the synergic articulation of brain areas divided in three subnetworks that extend beyond the classical left-lateralized perisylvian language circuit. Critically, we isolated a subregion of the left angular gyrus showing a significant interaction between gender congruency and gender type. The functional interplay between the angular gyrus and left perisylvian language-specific circuit proves crucial for constructing coherent and meaningful messages. Importantly, using graph theory we show that this complex system is functionally malleable: the role each node plays within the network changes depending on the available linguistic cues.

**Significance Statement:** Neural networks can be described as graphs comprising distributed and interconnected nodes. These nodes share functional properties but also differ in terms of functional specialization and the number of interconnections mediating the efficient transfer of information. Previous work has shown functional connectivity differences based on graph-theory properties between typical and atypical samples. However, here we have used concepts from graph theory to characterize connectivity during language processing using task-related fMRI. This approach allowed us to demonstrate how linguistic input drives brain network configuration during language comprehension. This is the first evidence of functional flexibility in language networks: the communicative capacity of each hub changes depending on whether the linguistic input grants access to meaning or is purely formal.

## Introduction

Independently of the structural diversity across human languages, their basic core falls on the need to transfer information between individuals (e.g., ideas, concepts, intentions, etc.) following some specific rules. Mastering the rules and the mechanisms that control the way in which these rules operate to structure words into phrases and sentences is essential for language comprehension. However, linguistic signals must also be anchored to conceptual knowledge in order to map linguistic inputs to their referents in the external world. It is, therefore, reasonable to infer that the interplay between form and meaning is fundamental for language comprehension. However, despite the significant increase in the knowledge of language processing, it is still not clear whether, how, and where conceptual information mediates the building of syntactic structures (1-8). The vast majority of studies have explored formal and conceptual factors separately, assuming there is no interaction between them (4, 8-11). The current study is aimed at shedding light on this issue by pinpointing how the neural network(s) underlying the building of syntactic structures combines the formal and conceptual factors embedded in our linguistic code.

Despite considerable research and both technological and methodological advances in noninvasive neuroimaging techniques over the last few years, how the language system process and integrates different types of formal and conceptual information remains controversial. Assessing the synchrony and interplay between distributed neural regions might be key to understanding how the language system operates. However, to date, no available fMRI study has characterized the networks underpinning formal and conceptual combinatorial mechanisms using both patterns of activation (i.e., mapping functional sensitivity and specificity across conditions) and connectivity profiles (i.e., tracking the interconnections that mediate the flow of information) without applying region-based theoretical constraints (12, 13). The use of network-based approaches and in particular graph theory to fMRI measures offers a unique opportunity to understand such a complex and multifaceted process as language.

According to graph theory, neural networks can be described as complex graphs comprising a certain number of distributed and interconnected nodes (14, 15). These nodes share functional properties but also differ in terms of both functional specialization (i.e., modularity, clustering, localm efficiency, etc.) and numbers of interconnections mediating the efficient transfer of information (i.e., centrality, links per node, node strength, etc.). These properties are used to define the node functional profile: by combining these indexes, it is possible to identify so-called hubs, regions with high communication capacity that are crucial for language functions (14, 15). By taking advantage of this network-based approach, we can address new questions in the field of language cognition: *Can the language network re-orchestrate the function(s) of critical nodes depending on the available linguistic information?*

Current neuro-anatomical accounts of language processing propose that the language system is a large-scale interactive set of networks (16-18). Each one of these networks is composed of distributed grey matter regions which are functionally related through long-and short-distance white matter fiber tracts (13). More specifically, it has been suggested that functional interaction between left-lateralized temporal, frontal, and parietal brain areas is crucial for all language-related operations (16, 17). Each one of these gray matter regions, located around the Sylvian fissure, has been functionally characterized combining cytoarchitectonical and functional criteria. According to current models, these regions are organized following a functional division into two main streams that reflect a clear segregation between syntax and semantics: mapping between form/sound and meaning is subserved by various white matter fiber tracks (the inferior longitudinal fasciculus, inferior fronto-occipital fasciculus, uncinate fasciculus, and extreme capsule), which connect the middle/inferior temporal gyri and the ventral part (pars orbitalis) of the inferior frontal gyrus (IFG) along the ventral stream; mapping from form/sound to syntax and articulation is subserved by the arcuate fasciculus, which connects the superior temporal gyrus and the dorsal part (pars opercularis) of the IFG along the dorsal stream (13, 16, 17, 19). Although these models have provided us with invaluable insights for understanding the neurobiology of language, the recent technological and methodological advances in neuroimaging analysis offer a unique opportunity to unravel the architectonic configuration of the language system by changing the focus from the actors to the dynamic interactions established between them.

The Spanish gender agreement system offers a convenient tool for experimentally isolating formal and conceptual linguistic information. Gender agreement systems comprise procedural mechanisms concerning the regular assignment of gender feature values (e.g., feminine, masculine) associated with sentence constituents. In Spanish, gender, in most cases, is signaled in a redundant way: determiners, adjectives, and pronouns must agree with noun gender, in both local as well as non-local relations. Nouns can also be distinguished in terms of whether gender carries semantic or strictly formal information. Most nouns referring to animate entities are assigned to masculine or feminine gender according to the biological sex of the referent (the conceptual system, according to 20). However, gender assignment for nouns that refer to inanimate entities or abstract concepts without biological sex, follows morphological criteria, such as whether the noun ends in “–a” or “–o”: Nouns ending in “–o” are usually masculine, while nouns ending in “–a” are usually feminine (the formal system, according to 20 with conceptual gender nouns also following this regularity in most of the cases (abuelomasc./abuelafem. [grandfather/grandmother]). Capitalizing on these linguistic properties characterizing the Spanish gender system, we were able to experimentally disentangle formal and conceptual factors by manipulating gender congruency between nouns and other parts of speech as well as noun gender type – i.e., whether nouns had *formal* or *conceptual gender*. This design enabled us to isolate purely syntactic mechanisms from those where an interplay of conceptual and formal factors is critical to establishing syntactic relations. In the current study, we conducted an fMRI experiment in which participants were required to read and then classify sentences according to their grammaticality. We followed a 2×2 factorial design with Gender Congruency [*Gender Match* and *Gender Mismatch*] between subject nouns and predicative adjectives and Gender Type [*Conceptual* and *Formal*] as factors.

A previous fMRI study by our group investigated syntactic combinatorial processes in minimal linguistic relations such as transparent and opaque determiner-noun phrases (e.g., *transparent*: “la_fem.sing._ casa_fem.sing._”, [the house]; *opaque*: “la_fem.sing._ fuente_fem.sing._” [the dish]) (7). This study suggested that the interplay between formal (i.e., gender-to-ending regularities) and conceptual (i.e., lexico-semantic) cues is critical even when the two grammatical elements are morphologically marked, as in the case of transparent relations. This interplay was mapped to a left-lateralized network which included highly selective language-related regions, such as the posterior MTG/STG, the pars triangularis within the IFG, and the supramarginal gyrus, but also other heteromodal areas such as the angular gyrus (12, 21). Extrapolating from these results to high-level sentence contexts, we hypothesized that during language comprehension word meaning interacts with syntactic information. If so, we should find an interaction between Gender Congruency and Gender Type. The contrast between grammatical and ungrammatical constructions should give rise to different activation patterns depending on whether nouns have conceptual or strictly formal gender information. However, there is a large distance in terms of combinatorial operations between isolated determiner-noun pairs and high-level sentence context. Therefore, if conceptual and formal gender agreement systems rely on similar neural mechanisms, we would expect no interaction between Gender Congruency and Gender Type: the contrast between grammatical and ungrammatical constructions should give rise to an overlapping network regardless of noun type (i.e., conceptual or formal).

Despite some differences, all theories aiming to explain how combinatorial processing occurs agree that processing and integration of linguistic information during the online comprehension of phrases and sentences involves inferior frontal and inferior/middle/superior temporal regions (10, 16 for divergent point of views). However, importantly, differing from other models but in consonance with our previous findings, the Memory, Unification, and Control model (MUC, 9, 16) assumes that parietal areas play a critical role within this system and are functionally related to the classic language-specific frontotemporal network. Thus, whereas both parietal and temporal regions subserve memory-related operations allowing for the access and retrieval of various types of information, the left IFG is a central hub not specific to language that is responsible for unification operations. According to this view, there is a direct dependency between the type of information required to decode the input (i.e., morphological/phonological, lexical, semantic, or syntactic) and the specific nodes recruited within the language circuit. It is proposed that an anterior-ventral to posterior-dorsal functional gradient similarly applies to frontal, temporal, and parietal clusters: while ventral areas contribute to lexico-semantic processing, dorsal areas are involved in syntactic operations (16).

Here, we leverage the synchrony between this model and our previous results to anchor our specific predictions (7). If the interplay between the left-lateralized frontotemporal network and parietal areas such as the left angular gyrus is critical for combining different information sources, we expected to find a neural trace of this functional relation across all conditions. However, we also expected variations in the activation and/or connectivity profiles of parietal areas as a function of the availability of semantic gender information – i.e., *conceptual* or *formal gender* (see 22 for convergent MEG results). Consistent with our previous findings, we expected that the dynamics and topology of the left angular gyrus would be affected by conceptual gender due to the increased combinatorial effort involved in integrating formal and conceptual information (6, 7). Critically, the interplay between formal and conceptual cues is not foreseen in the currently available models.

## Results

### Behavioral results

Mean decision times and error rates for each condition are presented in Table 1, with the corresponding standard error in parentheses. The percentage of correct responses was above 96 % for all experimental conditions, indicating that participants judged the well-formed sentences to be grammatically acceptable in contrast to the sentences with a gender agreement violation. 2×2 ANOVAs on mean decision times and error rates were performed using Gender Type (*Conceptual* and *Formal*) and Gender Congruency (*Match* and *Mismatch*) as factors. These analyses showed no significant effects among the experimental conditions, neither for accuracy nor for decision times (p > 0.05).

**Table 1.** Error rates (in %) and mean decision times (in ms) for conceptual and grammatical gender in the two types of sentences (match and mismatch) with standard error between parenthesis.

|  | Mean decision times |  | Error rates |  |
| --- | --- | --- | --- | --- |
|  | Match | Mismatch | Match | Mismatch |
| **Formal** | 680.19 (28.40) | 722.60 (24.66) | 3.94 (0.75) | 4.02 (1.04) |
| **Conceptual** | 688.44 (29.62) | 665.32 (28.40) | 3.72 (0.80) | 5.27 (1.26) |

### GLM-based activation analysis: Congruency effect (Gender Match vs. Gender Mismatch)

The main effect of *Congruency* included regions with higher responses for the *Gender Mismatch* than for the *Gender Match* condition and regions that exhibited the opposite pattern, i.e. higher activation for *Gender Match* than for *Gender Mismatch* condition. On the one hand, significant activation increases emerged from the *Gender Mismatch* > *Gender Match* contrast, including regions such as the right and left insula, the left pars orbitalis, opercularis, and triangularis within the IFG, the left precentral, left supplementary motor areas, and the left inferior parietal gyrus. On the other hand, the contrast *Gender Match* > *Gender Mismatch* produced higher brain response in regions such as the angular gyrus, anterior cingulate cortex, precuneus, middle temporal gyrus (MTG) and orbitofrontal cortex bilaterally as well as the left occipital and left superior frontal cortex (see Table 2 for a detailed list of regions; see Figures 1 for a representation of response patterns [activation or de-activation with respect to the fixation baseline]). Note that the hemodynamic response functions of these regions exhibited different patterns. The left anterior cingulate cortex, left middle temporal gyrus and left pars opercularis – within the IFG –, exhibited significant congruency effects between 2.5 and 5.0 seconds, whereas others regions such as the pars triangularis within the IFG, middle frontal gyrus and medial superior frontal gyrus exhibited sustained responses – i.e., for between 2.5 and 12.5 seconds – with significant differences in response amplitude between congruent and incongruent items (for more details see Figure 1).

**Table 2.** Significant activation clusters resulting from the main effect of Congruency (Match > Mismatch and Miamatch > Match) for conceptual and grammatical gender.

| Contrast | Hemisp | Region | x,y,z {mm} | Peak level | Cluster level |
| --- | --- | --- | --- | --- | --- |
|  |  |  |  | T | No. Vox |
| Mismatch > Match | L | Inferior Parietal Gyrus | -58 -46 38 | 4.35 | **1562** |
|  |  | Supramarginal Gyrus | -52 -46 38 | 3.88 |  |
|  |  | Angular Gyrus | -33 -50 38 | 3.13 |  |
|  |  | Middle Frontal Gyrus | -40 50 2 | 3.81 | **2384** |
|  |  | Pars Triangularis (IFG) | -52 20 30 | 3.75 |  |
|  |  | Pars Orbitalis (IFG) | -38 46 0 | 3.72 |  |
|  |  | Pars Opercularis (IFG) | -54 8 24 | 3.54 |  |
|  | R | Inferior Parietal Gyrus | 50 -34 58 | **5.23** | **2717** |
|  |  | Angular Gyrus | 40 -48 40 | 3.79 |  |
|  |  | Supramarginal Gyrus | 54 -38 42 | 3.42 |  |
|  |  | Precentral Gyrus | 38 -12 58 | 4.24 |  |
|  |  | Postcentral Gyrus | 28 -36 68 | 4.06 |  |
| Match > Mismatch | L | Superior Frontal | -12 54 36 | 3.14 | <u>83</u> |
|  |  | Orbitofrontal Medial | -4 46 -12 | 4.23 | **633** |
|  |  | Superior Frontal Medial | -6 52 6 | 2.49 |  |
|  |  | Precuneus | -6 -56 14 | 4.35 | **867** |
|  |  | Middle Occipital | -44 -74 34 | 3.22 | <u>167</u> |
|  |  | Inferior Occipital | -26 -94 -8 | 4.79 | **651** |
|  |  | Middle Temporal Pole | -52 10 -26 | 2.92 | <u>62</u> |
|  |  | Superior Temporal Pole | -42 20 -24 | 2.83 |  |
|  |  | Middle Temporal Gyrus | -56 2 -26 | 2.68 |  |
|  | R | Superior Frontal Medial | 2 56 16 | 2.69 | <u>83</u> |
|  |  | Anterior Cingulate | 8 48 24 | 2.93 |  |
|  |  | Middle Temporal Gyrus | 52 0 -24 | 2.91 | <u>214</u> |
|  |  | Precuneus | 6 -56 18 | 3.42 | **867** |
|  |  | Calcarine | 22 -96 -4 | 3.75 | **699** |
|  |  | Inferior Occipital | 32 -88 -8 | 3.53 |  |
|  |  | Middle Occipital | 34 -86 12 | 3.13 |  |

Only those clusters with an effect corrected with FWE or FDR criteria were considered as significant and it was included in the table. x,y,z {mm}= Coordinates in MNI space of local maxima. T = T scores. No.Vox. = Number of voxels significantly activated inside the cluster belonging to each local maximum. T scores at the peak level are reported in bold if they are significant after FWE correction.

**Figure 1.**
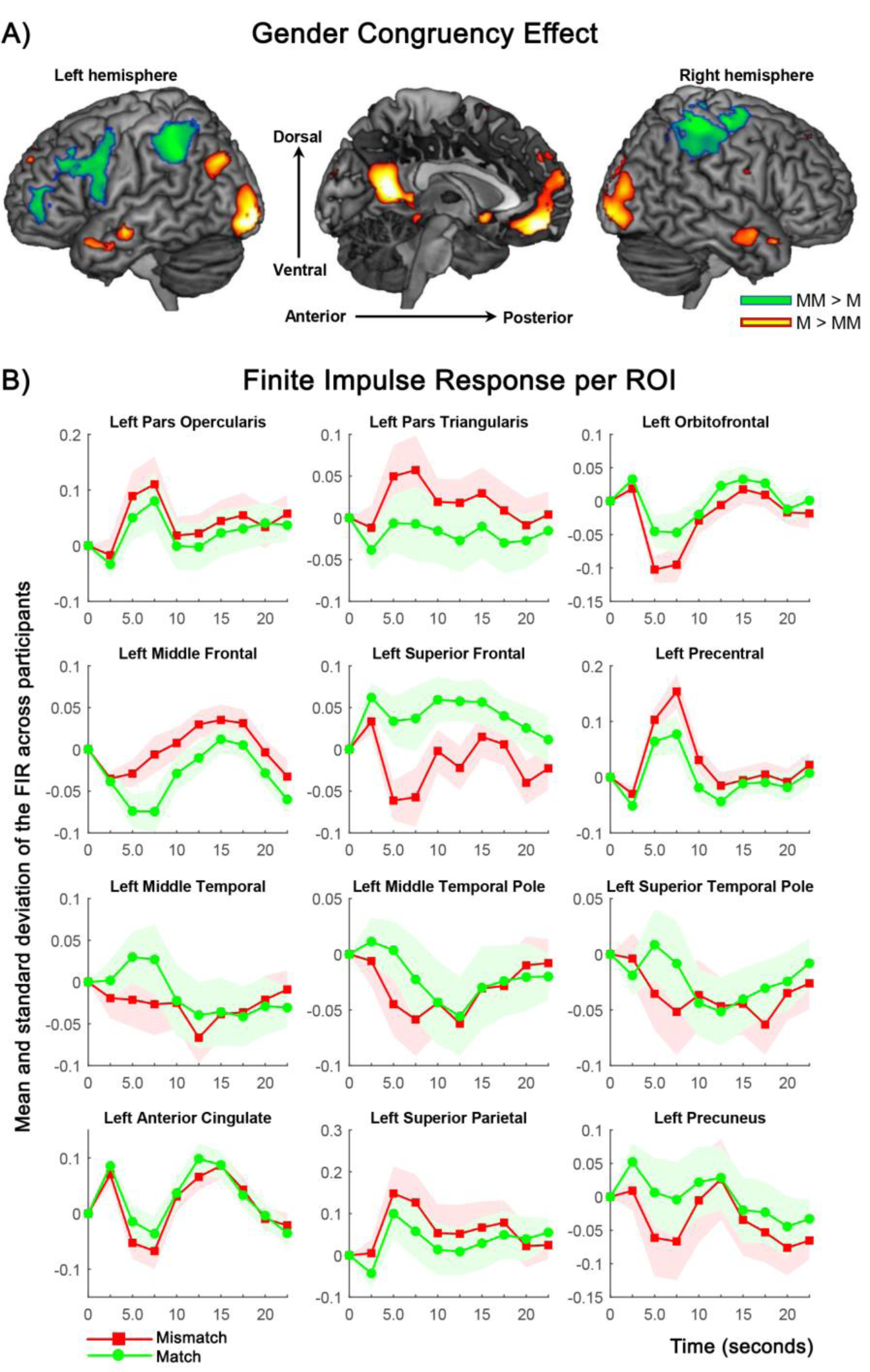
GLM-based activation analysis: Congruency effect. (A) Statistical parametric map emerging from the main effect of Gender Congruency was projected on the MNI single-subject T1 image. The two tails of the t-contrast were represented with different colors: *Gender Mismatch* > *Gender Match* in green and *Gender Match* > *Gender Mismatch* in red-yellow. All clusters depicted were statistically significant with a p-value corrected for multiple comparisons. MM: Gender Mismatch; M: Gender Match. (B) HRFs of those regions resulting from the main effect of Gender Congruency. Gender Mismatch and Gender Match are represented in different colors. The vertical dotted lines depict the maximum amplitude peaks and the latency corresponding to the location of this maximum.

### GLM-based data analysis: Interaction between Gender Congruency and Gender Type

We observed significant interaction effects in two different clusters including the left inferior parietal gyrus and the left angular gyrus (Figure 2 and Table 3). Significant differential effects emerging from the contrast *Mismatch* vs. *Match* in both regions were found only for *Conceptual Gender*. No significant differential effect was found in these regions for *Formal Gender*. Note that the hemodynamic response functions of these two parietal clusters were very similar; both exhibited an increment in the response pattern between 2.5 and 7.5 seconds with a maximum peak around 5.0 seconds.

**Table 3.** Interaction between Conguency Pattern (Match and Mismatch) and Type of Gender (Conceptual and Formal).

| Region | Interaction |  |  | Match vs. Mismatch for Conceptual Gender |  |  |
| --- | --- | --- | --- | --- | --- | --- |
|  | x,y,z {mm} | T | No. Vox. | x,y,z {mm} | T | No. Vox. |
| Inferior Parietal (L) | -46 -56 48 | 4.12 | **998** | -56 -46 40 | **5.76** | **506** |
| Angular Gyrus (L) | -40 -62 40 | 3.05 |  | -42 -54 44 | **4.57** | **65** |

Only those clusters with a significant interaction effect ( $p < 0.001$ uncorrected) were reported. x,y,z {mm}= Coordinates in MNI space of local maxima. Vx = Number of voxels significantly activated inside the cluster belonging to each local maximum. T = T scores. T scores are reported in bold if they are significant at the peak level after FWE or FDR correction.

**Figure 2.**
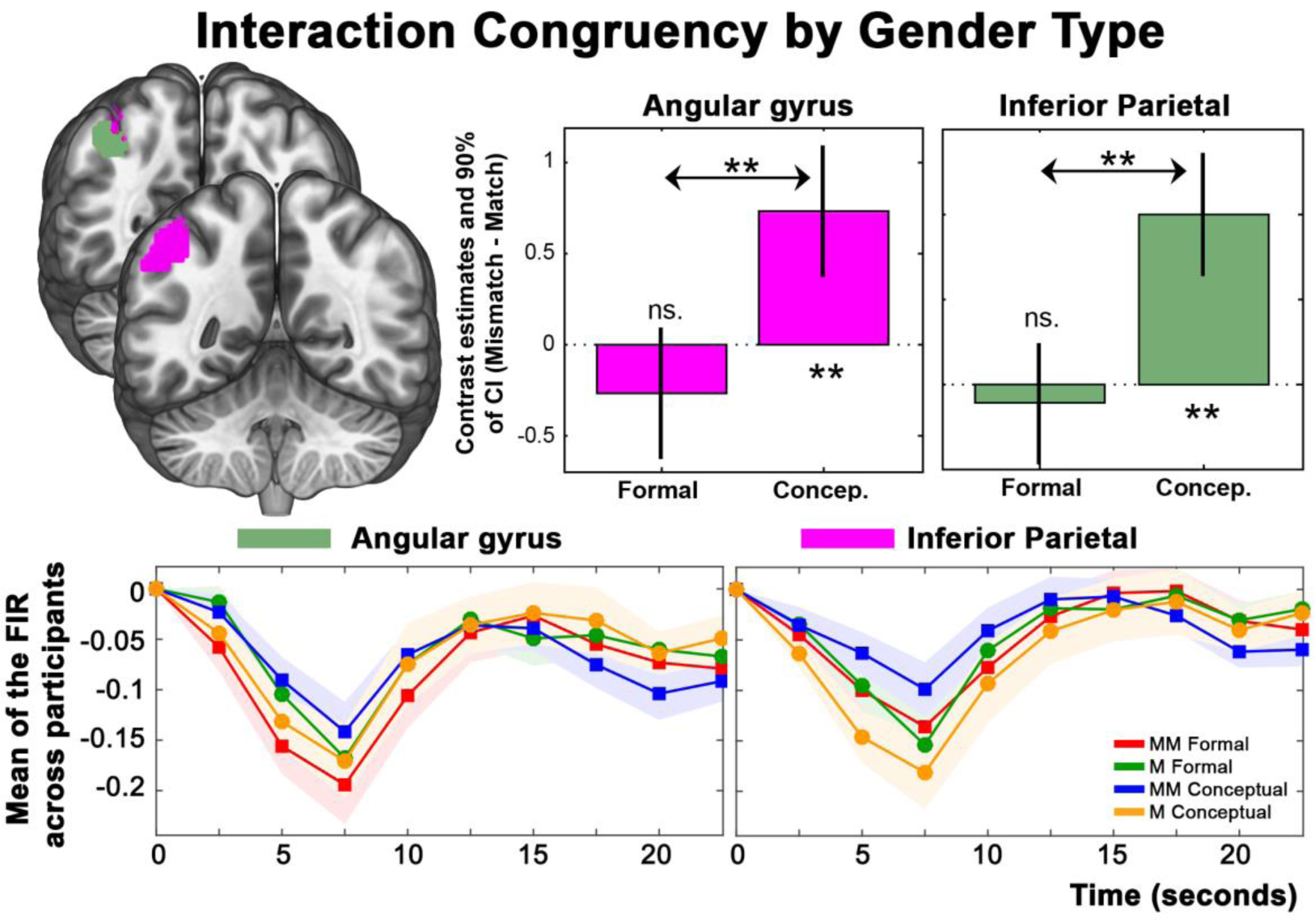
GLM-based activation analysis: Interaction effect. Statistical parametric maps emerging from the interaction effects between Gender Congruency and Gender Type were projected on the MNI single-subject T1 image. The sagittal and axial views represented in the upper part of the figure display the significantly activated clusters. The bar graphs display the contrast estimates and 90% of confidence intervals at the maximum peaks representative of the two clusters resulting from the interaction effect. Time courses of the HRFs are represented in the lowest part of the figure. Each condition is represented in different colors. The vertical dotted lines signal the maximum amplitude peaks and the latency corresponding to the location of this maximum. MM: Mismatch; M: Match. CG: Conceptual Gender; FG: Formal Gender.

**Figure 3.**
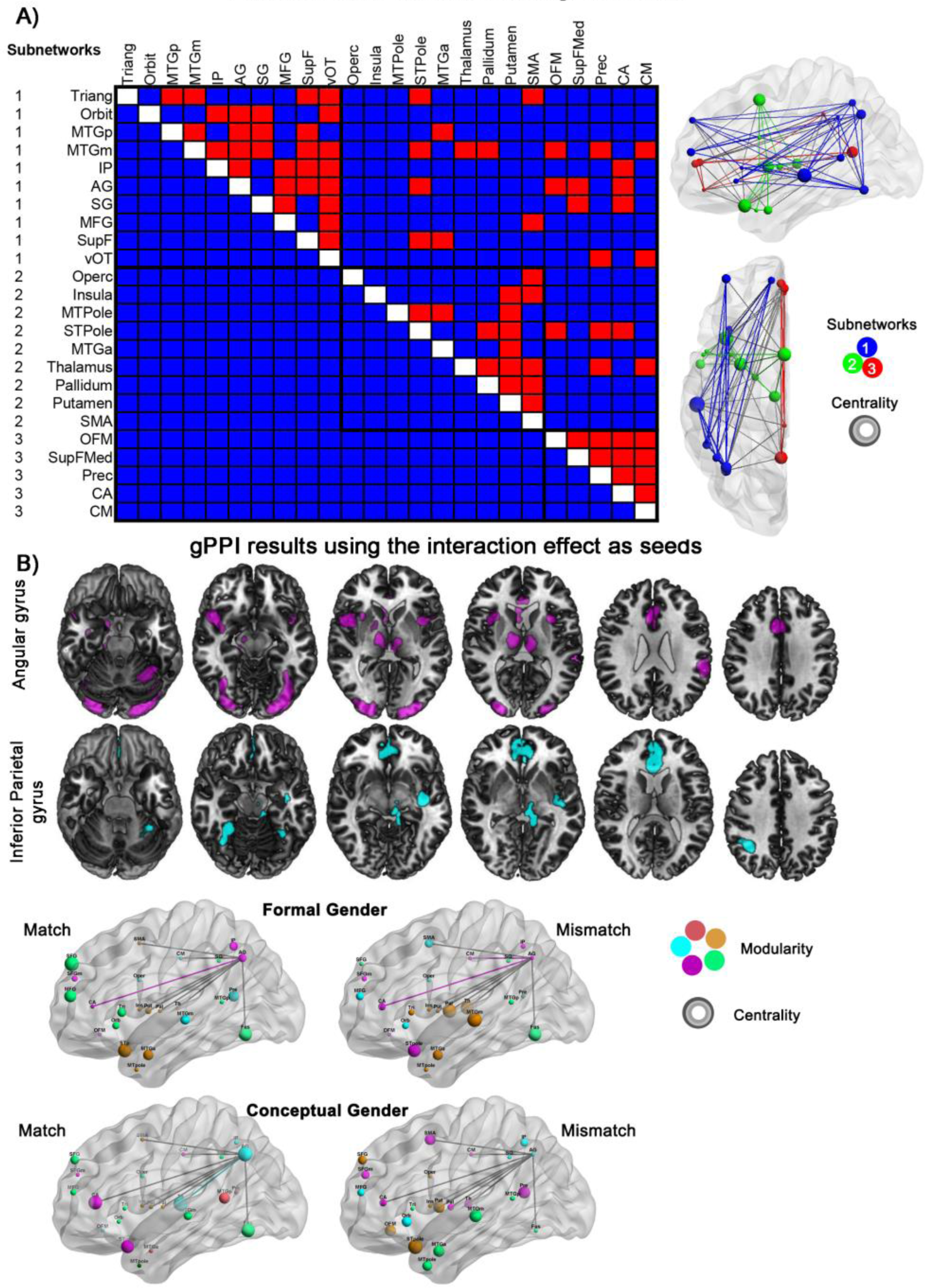
Changes in brain network configuration as a function of the linguistic input properties. (A) Matrix of functional interactions between the 24 regions of interest. Red indicates a significant connection between two areas, blue no connection. Values on the left (between 1 and 3) indicate the subdivision of the network into non-overlapping groups of nodes. In the right part, all the significant interactions are superimposed in a 3D reconstruction of a brain surface template (48, http://www.nitrc.org/projects/bnv/bnv/). The three subnetworks are represented in different colors. (B) Significant second-level group effects resulting from the gPPI using the angular gyrus and the inferior parietal gyrus as seeds. (C) Individual changes in modularity and betweeness centrality across experimental manipulations superimposed in a 3D reconstruction of a brain surface template. Modularity and centrality are displayed by the color and the size of the spheres respectively.

### Connectivity results

In order to characterize the synchrony and interplay between the 24 nodes resulting from the group-level F-test contrast *All conditions vs. Null*, condition-dependent functional connectivity analyses were performed at both global and local network scales. Firstly, we used the Network-based Statistic Toolbox (NBS, 23) to compare the global network configuration across conditions. These analyses confirm what was shown by the FIR models and extend this result revealing interesting details about the functional connectivity profile of each node. Significant direct interactions were found in 79 out of 276 possible connections (see Figure 4A). This analysis reveals three functionally different sub-networks. Each subnetwork included short-and long-distance connections. Subnetwork 1 covered frontal, temporal and parietal areas: middle and superior frontal gyri, pars triangularis, and orbitalis within the IFG, posterior and medial part of the MTG, inferior parietal and angular gyri, supramarginal gyrus, and fusiform gyrus. Subnetwork 2 comprised pars opercularis within the IFG, insula, the most anterior part of the MTG/STG including the temporal pole, premotor regions, and basal ganglia (i.e., thalamus, pallidum, and putamen). Subnetwork 3 included orbitofrontal areas, the medial part of the superior frontal gyrus, precuneus, and middle and anterior cingulate cortex. Figure 4A contains a detailed characterization of the connectivity profile of each node. Focusing on the linked nodes, there were no differences between experimental conditions: A single network configuration underlies the processing of conceptual and formal information during sentence comprehension (see Figure 4A).

**Figure 4.**
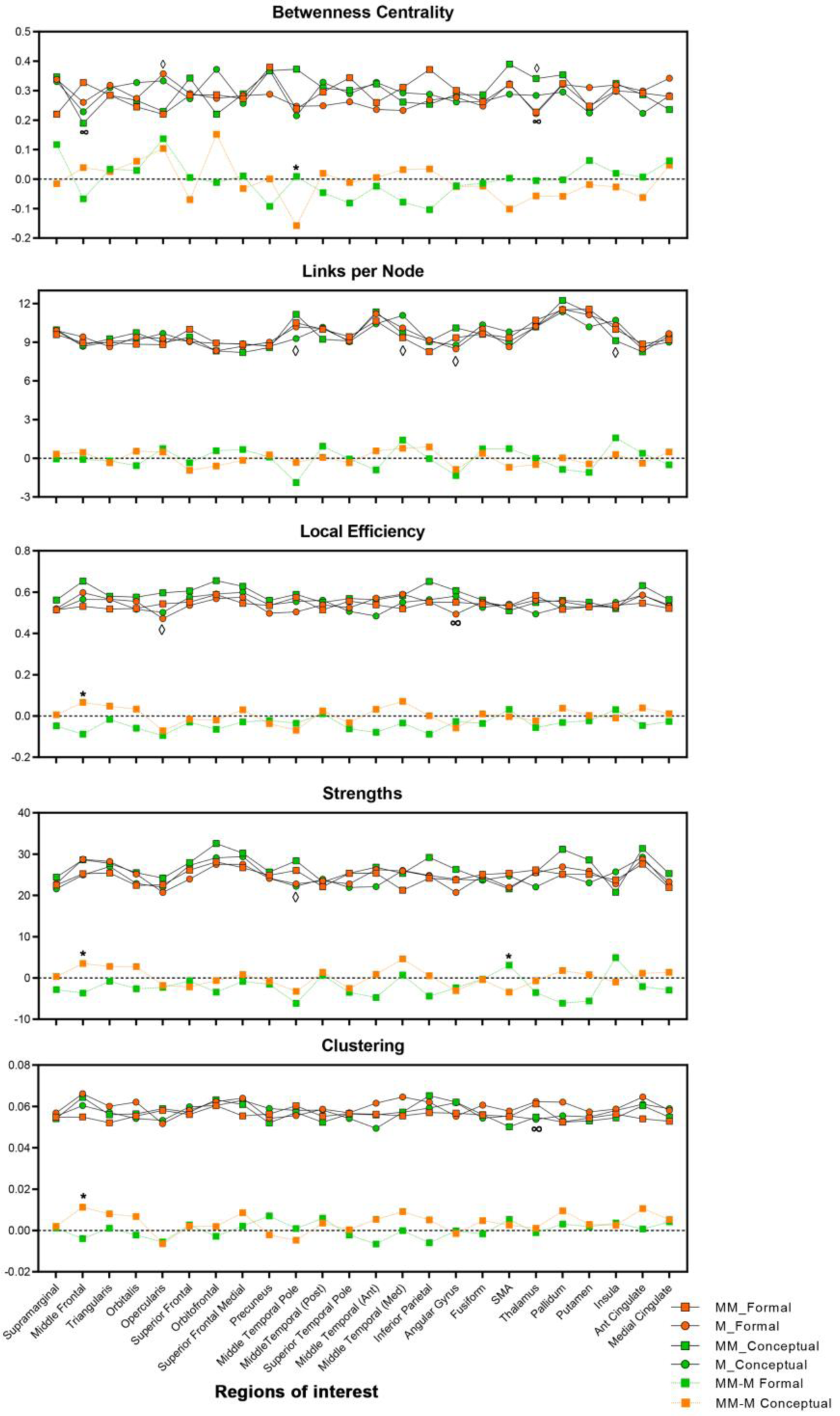
Topological characterization of the network as a function of the experimental manipulation. Each box represents the change in a graph-theory property across nodes. Five different measures were included: 1) node betweenness centrality; 2) node strength; 3) links per node; 4) local efficiency; 5) weighted clustering coefficient. Significant main effects and interactions are signed with one of these symbols (∞ for Gender Type, ? for Grammaticality, * for the interaction between Gender Type and Grammaticality). The upper part of each graph comprises the average value over the participants per condition, while the lower part represents the interaction between Gender Type and Grammaticality. Each symbol/line in the lower part represents the difference between the Mismatch and the Match conditions.

Secondly, we specifically explore the whole-brain connectivity profiles of the angular and the inferior parietal gyri, the two regions exhibiting significant interaction effects (i.e., *Gender Congruency x Gender Type*). Angular gyrus response was tightly coupled with activity in the pars orbitalis within the IFG, middle and superior frontal gyri, orbitofrontal areas, the posterior and medial parts of MTG, superior temporal pole, fusiform gyrus, and anterior cingulate cortex, whereas only a subset of these areas (i.e., middle and superior frontal gyri, the medial part of MTG, fusiform gyrus, and anterior cingulate cortex) showed significant connections with the inferior parietal gyrus (see Figure 4B, 4C, and Table 5 for the whole-brain second level gPPI results of these two critical ROIs).

**Table 5.**
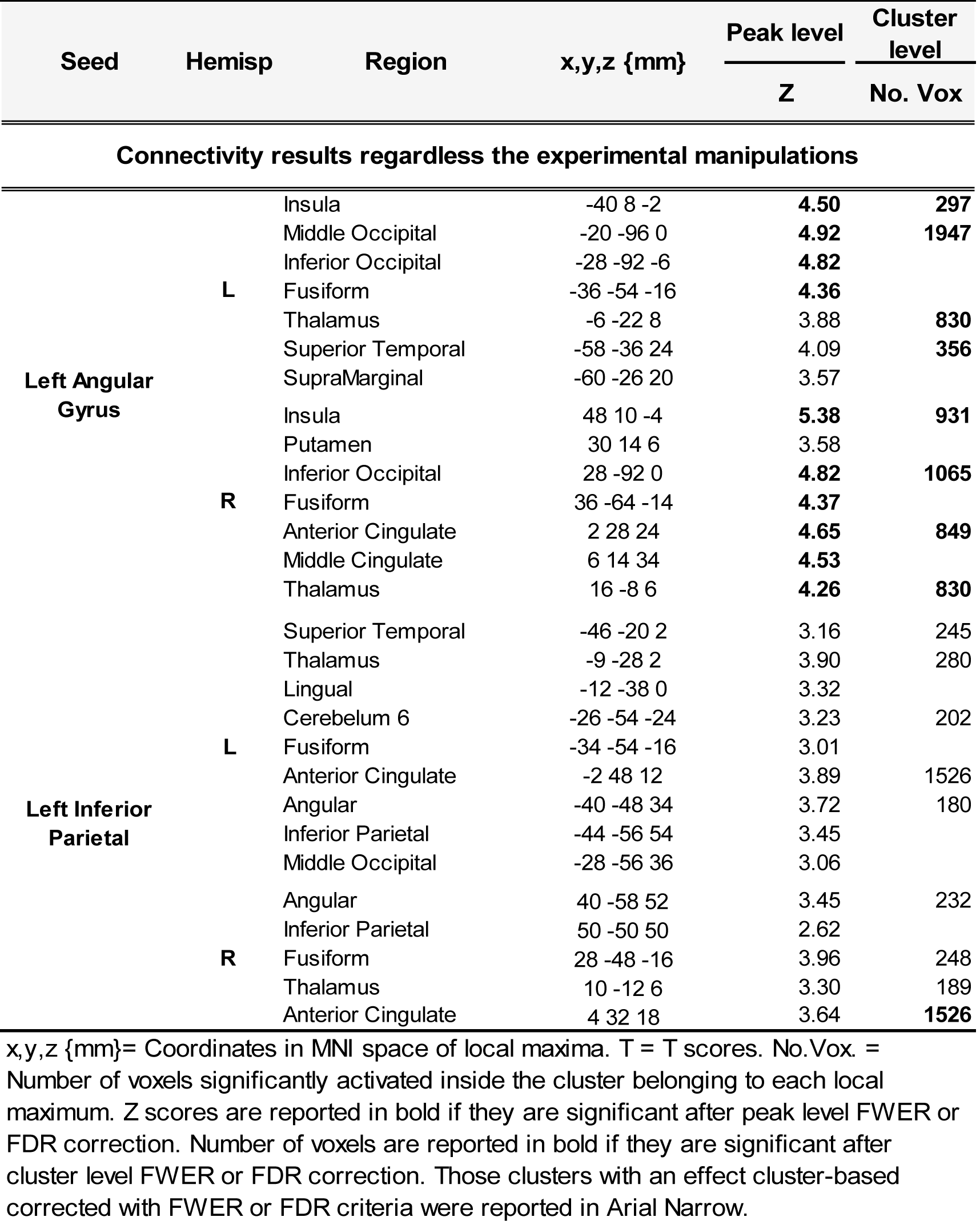
Whole-brain connectivity results using the two ROIs resulted from the the interaction between Conguency Pattern (Match and Mismatch) and Type of Gender (Conceptual and Formal) as seeds.

Thirdly, we described how the topology of the network changes as a function of our experimental manipulation using six graph-theory measures: 1) node betweenness centrality; 2) node strength; 3) links per node; 4) local efficiency; 5) weighted clustering coefficient; 6) modularity. Based on these measures, we demonstrated how the communicative capacity of each hub changes depending on the available linguistic cues. On the one hand, the middle temporal pole, the medial part of the MTG, the insula, the angular gyrus, the thalamus, and the pars opercularis of the IFG were identified as critical hubs for processing grammatical relations, while the thalamus, the angular gyrus, and the middle frontal gyrus were pinpointed as critical interconnected nodes for decoding gender-related information. On the other hand, the middle temporal pole, the premotor regions, and the middle frontal gyrus emerged as critical interconnected hubs mediating the integration of formal and conceptual information (see Figure 4C for the representation of one prototypical participant). In Figure 5, each graph-theory property is represented across regions and experimental conditions. Surprisingly, this approach demonstrated that all the 24 nodes we included in our analyses similarly contribute to the processing of conceptual and formal information during sentence comprehension. The distribution of the graph properties was very similar across the 24 nodes. To explore the integration capacity of the network across conditions, we computed the optimal community structure at both global (i.e., maximized modularity) and local scales (i.e., a subdivision of the network into non-overlapping groups of nodes). In Figure 1S, the maximized modularity was presented for each experimental manipulation. As shown, there were no significant differences across conditions.

**Figure 5.**
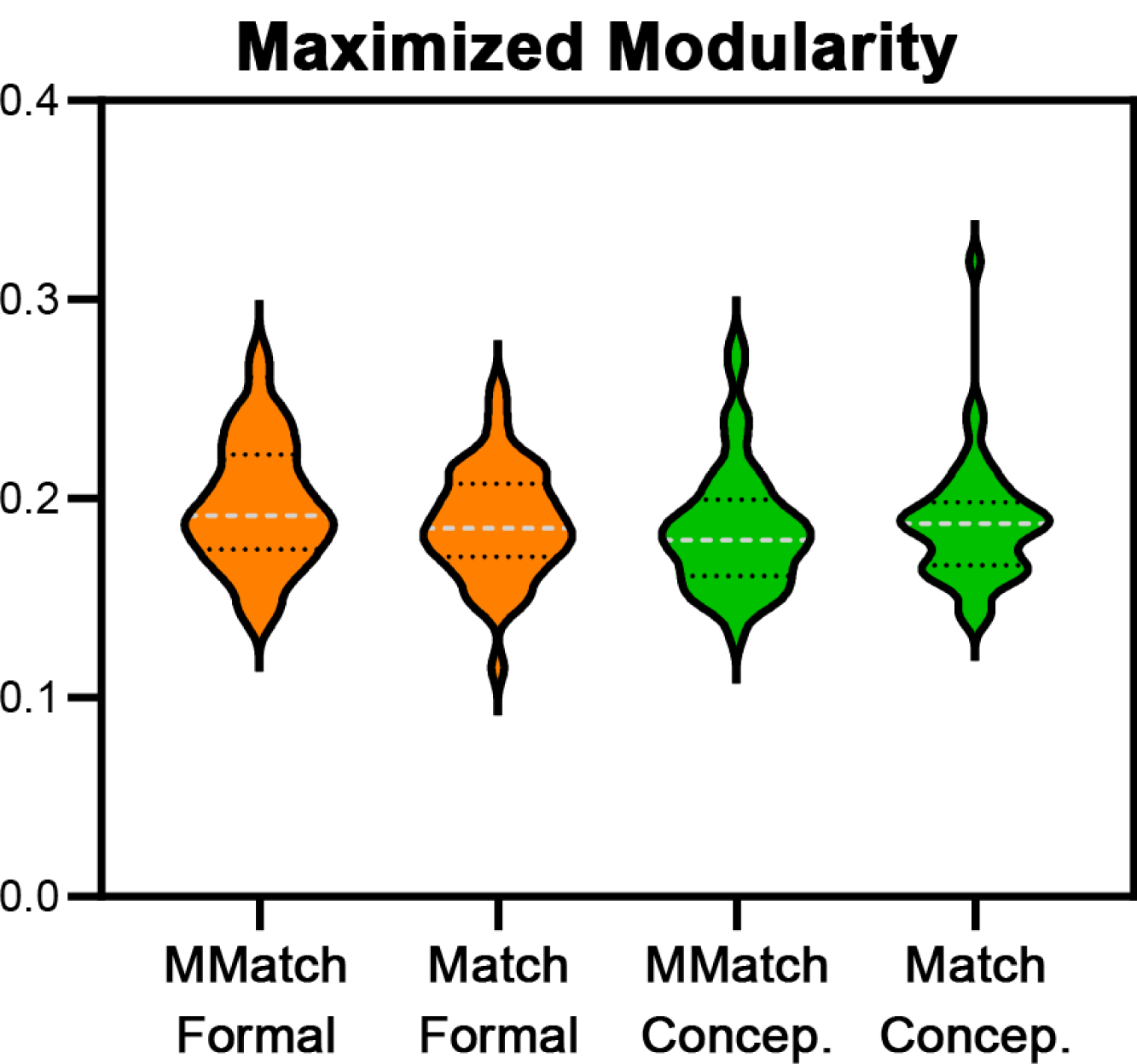
Optimal community structure at the global scale (i.e., maximized modularity (Q), see Brain Connectivity Toolbox for more details). The distribution of the maximized modularity per experimental manipulation is depicted using the violin plots. The dotted grey line represents the median per condition.

## Discussion

The present fMRI study demonstrated how multiple neural mechanisms operate in a coordinated fashion during the processing and integration of the formal and conceptual information required for comprehending grammatical relations. In line with previous studies, this system involves a left-lateralized perisylvian circuit typically associated with language-specific functions (10, 16, 17). However, our data suggest that bilaterally distributed domain-general regions, comprising cortical and subcortical areas, are concurrently engaged during this process. We described how this system comprises three functionally different subnetworks. While subnetworks 1 and 2 comprised regions previously related with language functions (e.g., IFG, MTG, inferior parietal, supramarginal and angular gyri, and fusiform gyrus), subnetwork 3 included areas previously linked with general attentional control mechanisms (e.g., orbitofrontal, superior frontal gyrus, precuneus, and middle and anterior cingulate cortex). Despite this functional segregation, we found clear evidence for interactions between them. However, the main contribution of this study was the parietal (i.e., angular and inferior parietal gyri) involvement during the integration of formal and conceptual information associated with the different grammatical elements. The amount of connections between these regions and the classical left frontotemporal language network (i.e., subnetwork 1) points out the critical role this parietal region must play as part of this system. Graph-theory properties characterizing the angular gyrus display its functional malleability: the role the angular gyrus plays within this network changes depending on the available linguistic cues.

Our data indicate that interaction between formal and conceptual cues, rather than serial processing of these two types of information, is essential for the proper interpretation of grammatical relations. Combining these two dimensions – i.e., conceptual and formal features – within the same linguistic representation constitutes a new fMRI approach that allowed us to draw stronger conclusions. Using this type of manipulation, it was possible to map formal and conceptual integration processes without confounding mechanisms triggered by the computation of ungrammatical constructions. The network-based analyses performed here confirm what was shown by the activation patterns and the FIR time courses and reveal new interesting details about the functional connectivity profile of each node. Critically, this approach allowed us to demonstrate how the interface between formal and conceptual features depend on the synergic communication of brain areas extended beyond the classical left-lateralized language network and how this system is flexible enough to re-allocate the functional weight of its different nodes in response to specific input properties.

As we pointed out in the introduction, there is an extensive knowledge of the left-lateralized perisylvian circuit underlying the different language-related operations. The functional role played by the different nodes within this circuit in comprehension has been extensively explored (8, 11, 16). Specifically, there are numerous studies using typical and atypical populations focusing on the functional characterization of the temporal lobe – i.e., from anterior to posterior and from superior to inferior – and the inferior frontal gyrus (8-10, 16, 17). Even though these two brain regions have been the focus of these studies, it is quite clear that the language-related functions are supported by a complex system extending beyond this left-lateralized perisylvian circuit. Some of the most recent neuro-cognitive accounts tackling the neural dynamics underlying language comprehension reflect this complexity by taking into account other frontal and parietal nodes and its functional and anatomical connections with the well-established left-lateralized perisylvian circuit (9, 16, 18). However, most of the available functional and connectivity measures used to characterize the language network arise from a region-based approach (12). Despite its statistical strengths, this methodological approach has several weaknesses with a direct impact on the understanding of the language network(s) topology and dynamics.

The approach used in the current study reveals that the interplay between this left-lateralized perisylvian circuit and other cortical (e.g., angular and inferior parietal gyri, anterior cingulate cortex, superior and middle frontal gyri, etc.) and sub-cortical (e.g., thalamus, dorsal striatum, pallidum) regions is critical for the formal and conceptual combinatorial mechanisms started-up during agreement computation. The response patterns of some of these regions varied as a function of the congruency between different grammatical elements suggesting that grammatical violations imply additional processing costs in comparison with the integration of congruent information (5-7, 24). Even when the analysis of the activation profiles targets some nodes within the circuit as critical for certain experimental manipulation, all the functionally active nodes should play a role within this system and, thus, we explore the relationships established between them. This network-based approach enabled us to characterize this neural circuit providing chronological and topological details. Based on those analyses, we demonstrated for the first time the capacity this system has to handle the functional role of each node depending on the input properties. The fine-tuning of this system seems to be constrained by available conceptual and/or formal information. This functional malleability is supported by the redundant connectivity profiles characterizing the different nodes within this system.

Critically, our data provide clear findings indicating the crucial role played by the angular gyrus and adjacent inferior parietal areas during the integration of formal and conceptual features. The activation and connectivity profile of this area is affected by both *Gender Congruency* and *Gender Type*. This parietal area exhibited greater responses for incongruent than for congruent items. This difference was larger for *Conceptual* than *for Formal Gender* suggesting its sensitivity to conceptual, but also formal properties. Conceptual nouns appear to impose a processing cost in the integration of formal and conceptual information. The engagement of this area for the comprehension of grammatical relations points to an interface between a classical left-lateralized language-specific perisylvian circuit and a central hub in the highly heteromodal semantic associative system (7, 25-28).

The involvement of the angular gyrus during language-related operations is a fact underpinned by a large number of studies (see 16 for a revision of this topic). A variety of cognitive functions has been attributed to this parietal area including retrieval of different types of linguistic information – e.g., morphological, phonological, lexico-semantic, and/or syntactic information (16), syntactic (29), and semantic combinatorial operations (12, 25, 30). Trying to reconcile the heterogeneity of the results, Seghier (26) postulated that this area constitutes *“a cross-modal integrative hub that gives sense and meaning to an event within a contextualized environment, based on prior expectations and knowledge”* (25, 27, 28). In line with this proposal, the semantic-biased activation and connectivity profiles of this area presented here disclose its multifunctional role during the processing and integration of linguistic information. Top-down and bottom-up anatomical connections between the angular gyrus and multiple frontotemporal (e.g., the middle, superior and inferior frontal gyri, and the MTG) and medial (e.g., the hippocampus, caudate, and precuneus) regions underpin this proposal (see also 18, 26 for a review of this topic). Here we have shown, for the first time, how the activation and connectivity profiles of this parietal region support this hypothesis. The angular gyrus enjoys a close relationship with eleven out of 24 nodes, emerging as one of the critical interconnected hubs underlying the building of local grammatical relations. These connections cover regions spreading through all the subnetworks. The connectivity profile of this region comprises direct connections with the pars orbitalis within the IFG, the posterior and medial part of the MTG, the superior temporal pole, the fusiform gyrus, the middle and superior frontal gyri, the orbitofrontal cortex, the inferior parietal gyrus, and the anterior cingulate cortex (see 31). These interlinks were significant for all the experimental manipulations with no difference across conditions. The changes in local efficiency and links per node reported for the angular gyrus discloses its functional malleability: the role the angular gyrus plays within this network changes depending on the available linguistic cues.

As theoretical models of sentence processing have proposed, it is possible to distinguish two main streams that play different roles within the language-specific left-lateralized perisylvian circuit (10, 13). All of these models are consistent in proposing a clear segregation between syntax and semantics affecting the left IFG and the left MTG/STG as the critical hubs within this network. According to this dorsoventral functional distinction, we would have expected a lexico-semantic cost in ventral areas triggered by *Conceptual Gender* and a syntactic processing cost in dorsal areas associated with *Formal Gender*. However, results concerning brain activation and functional connectivity question this functional semantic-syntactic segregation: processing and integration of formal and conceptual information involve both ventral and dorsal streams (29, 32). The IFG and the MTG are redundantly and tightly connected to different cortical areas in both dorsal and ventral streams.

Specifically, pars orbitalis and triangularis within the IFG were tightly coupled to the response of different frontal and temporal nodes including superior frontal gyrus, superior and middle temporal poles, medial part of the MTG, fusiform gyrus and premotor areas. In contrast, the pars opercularis was connected exclusively to premotor areas. As far as the functional connectivity is concerned, it is likewise unclear the dual-route functional segregation. The anterior, as well as the posterior part of the MTG/STG exhibited direct connections with dorsal areas such as the angular and supramarginal gyri, but also ventral areas such as the pars triangularis and orbitalis within the IFG. These interlinks were significant for all the experimental manipulations with no differences across conditions. As has already been suggested, these connections are probably mediated by thalamo-cortical pathways, and/or basal ganglia loops, as it is shown by the activation and connectivity profile of the subcortical areas presented here.

The results presented here clearly question the classical language system comprising left-lateralized frontotemporal areas functionally segregated into two main streams. Language functions depend on a complex set of redundantly connected cortical and subcortical regions (sub-network 1 and 2) which transcend the limits of the classical left-lateralized perisylvian network. Parietal regions, but more importantly the functional interplay between parietal areas and the left perisylvian language-specific circuit was identified as crucial for constructing coherent and meaningful messages. Nodal changes on graph-theory measures across conditions suggest an intrinsic functional malleability affecting different nodes within this system. *Is the functional architecture of the language network similar across different subjects? Is there a causal link between the individual differences in language abilities and the functional architecture of the language network? Are the nodes within the language network equally malleable?* Further work should be directed (1) to explore the language network dynamics considering the functional malleability of each node; (2) to isolate the neuroanatomical route(s) through which these two systems communicate with each other; (3) to identify the switch that controls this interface; and (4) to characterize the functional architecture of the language network taken into account individual phenotypes.

## Supporting information

Figure 1S. Clustering indiviadual variability

## Materials and Methods

### Participants

Fifty-three healthy paid volunteers gave written informed consent to participate in this study. All were highly proficient speakers of Spanish and all gave informed consent as stipulated in the ethics approval procedure of the *BCBL Research Ethics Committee*. They all had right-handed dominance, normal or corrected to normal vision and no history of psychiatric, neurological disease or learning disabilities. The quality of the fMRI data of each individual was explored using the Artifact Repair toolbox (Gabrieli Cognitive NeuroScience Lab; http://cibsr.stanford.edu/tools/ArtRepair/ArtRepair.htm). After that, a total of forty-nine participants (thirty females), with ages ranging from 22 to 42 years (mean = 28.6, standard deviation = 4.8), were used to estimate group effects.

### Stimuli and experimental procedure

Each subject participated in a single functional run consisting of an event-related 2 × 2 factorial within-subject design. Each trial consisted of a word-by-word visual presentation of four-word sentences that were grammatically acceptable or not. Words were displayed for 300 ms in white capital letters on a black background. In order to optimize the sampling of the BOLD response, an interstimulus interval was included between successive sentences. During this period a fixation point (“+”) was presented for durations varying between 2 and 8 seconds across trials. This baseline period also allowed us to estimate the time course of the BOLD response associated with the critical word and to counteract possible expectation effects that might influence brain responses. After each sentence, a visual cue indicated that participants should distinguish whether the sentence was grammatically acceptable or not by pressing one of two buttons.

The stimuli consisted of 160 sentences, which included 120 critical items and 40 fillers. Each sentence contained a subject noun followed by a verb, which was always followed by a predicative adjective. The gender of the adjective (the critical word) was manipulated to produce agreement or disagreement with the subject noun. While half of the sentences included a gender agreement violation between the subject noun and the predicative adjective, the other half consisted of well-formed sentences. In addition, we also manipulated the gender type of the subject by including nouns with formal (e.g., *acera* [sidewalk]) and conceptual gender (e.g., *abuela* [grandmother]) in a 1:1 ratio. The resulting 2 × 2 factorial design comprised the Gender Type of the subject noun [*Formal Gender* and *Conceptual Gender*] and the Gender Congruency between subjects and predicative adjectives [*Gender Match* and *Gender Mismatch*] as factors (see examples (1) and (2) below).

1. **Formal Gender**
2. Gender Match: e.g., *La acera era* estrecha ([The sidewalk]_fem.sing._ was narrow _fem.sing._)
3. Gender Mismatch: e.g., \**La acera era* estrecho ([The sidewalk]_fem.sing._ was narrow _masc.sing._)
4. **Conceptual Gender**
5. Gender Match: e.g., *La abuela era* sabia ([The grandmother]_masc.sing._ was wise _masc.sing._)
6. Gender Mismatch: e.g., *\*La abuela era* sabio ([The grandmother]_masc.sing._ was wise _fem.sing._)

Depending on gender-to-ending regularities, nouns in Spanish can be distinguished in terms of whether they have transparent or opaque ending. The endings of *transparent nouns* have a regular correspondence with a specific gender class (“–a” for feminine and “–o” for masculine, e.g., libro_mas._ [book]; luna_fem._[moon]), while the endings of *opaque nouns* provide no information regarding the gender class to which the noun belongs (e.g., lápiz_masc._[pencil]; vejez_fem._[old age]). To avoid possible lexical and/or morphological confounds that could influence the appearance of an interaction effect, we controlled the gender predictability of conceptual and formal nouns by selecting nouns whose gender was morphologically marked by either “–a” or “–o” (i.e., *transparent nouns*). In order to prevent strategies related to the morphological decomposition of nouns and adjectives (i.e., participants only attending to suffixes in determining whether or not there was a gender grammatical violation), we included only opaque predicative adjectives, i.e. adjectives that end with non-canonical suffixes (e.g., “–e”, “–n”, “–l”) in filler sentences. In addition to gender agreement, in Spanish, it is mandatory that subject nouns and adjectives also agree in number. In order to avoid possible interactions between gender and number agreement features, number agreement was also controlled: a) all subject nouns and adjectives agreed in number; b) all subject nouns and adjectives were morphologically marked for number with the canonical Spanish plural suffix (“–s”) and c) half of the nouns were presented in singular form, and the other half in plural form. All nouns and adjectives were of medium lexical frequency [nouns: mean = 38.37 per million, SD = 54.25; adjectives: mean = 22.67 per million, SD = 61.65] and 4 to 9 letters long [nouns: mean = 5.69, SD = 0.91; adjectives: mean = 6.41, SD = 1.65] (see also Table 4) according to the Spanish ESPaL database (http://www.bcbl.eu/databases/espal/) (33).

**Table 4.** Mean of the frequency, length and neighbors of the critical word (predicative adjectives) per condition. Mean values for conceptual and formal gender in the two congruency patterns (match and mismatch) were included, with its corresponding standard deviation between parenthesis.

|  | Formal Gender |  | Conceptual Gender |  |
| --- | --- | --- | --- | --- |
|  | Mismatch | Match | Mismatch | Match |
| **Log (frequency)** | 2.2828 (1.33) | 2.5208 (1.54) | 1.6333 (1.84) | 1.9732 (1.72) |
| **Length** | 5.5667 (0.63) | 5.4333 (0.86) | 7.9231 (2.13) | 7.5385 (1.70) |
| **Lev Neighbors** | 1.5433 (0.25) | 1.5 (0.36) | 1.9981 (0.62) | 1.8538 (0.52) |

### MRI acquisition

The experiment was performed on a 3 Tesla Siemens MAGNETOM TIM Trio scanner, using a thirty two channel phased-array coil (Siemens, Erlangen, Germany). Functional event-related scans consisting of 544 echo-planar images were acquired using a T2*-weighted gradient-echo pulse sequence with the following parameters: field of view (FOV) = 192 mm; matrix = 64 × 64; echo time (TE) = 30 ms; repetition time (TR) = 2.5 s; flip angle = 90°. The volume was comprised of 32 axial slices with 3 × 3 x 3 mm^3^ voxels without slice gap. The first six volumes of each functional run were discarded to ensure steady-state tissue magnetization. In addition, a MPRAGE T1-weighted structural image (1 × 1 x 1 mm resolution) was acquired with the following parameters: TE = 2.97 ms, TR = 2530 ms, flip angle = 7° and FOV = 256 × 256 x 160 mm^3^.

### GLM-based activation analyses

Functional data were analyzed using SPM8 and related toolboxes (http://www.fil.ion.ucl.ac.uk/spm). Raw functional scans were slice-time corrected taking the middle slice as reference, spatially realigned, unwarped, coregistered with the anatomical T1 (34) and normalized to MNI space using the unified normalization segmentation procedure. Global effects were then removed using a global signal regression analysis (35), and after that the data were smoothed using an isotropic 8mm Gaussian kernel. Resulting time series from each voxel were high-pass filtered (128s cut-off period).

Statistical parametric maps were generated using a univariate general linear model with a regressor for each stimulus type obtained by convolving the canonical hemodynamic response function with delta functions at stimulus onsets, also including the six motion-correction parameters as regressors of non-interest. The stimuli onsets included five different components. The first corresponded to the onset of each sentence trial and was modeled as a single regressor, independently of the experimental conditions. The next four corresponded to the experimental conditions (*Formal Gender Mismatch, Formal Gender Match, Conceptual Gender Mismatch*, and *Conceptual Gender Match*) starting from the onset of the critical adjective. In addition, in order to control the intrinsic variability of the lexical variables, we also included three regressors with the parametric modulation of the canonical response according to the frequency, length, and Levenshtein’s distance of each item. Those trials associated with incorrect behavioral responses were removed from the corresponding condition and included in the model as an additional regressor of non-interest. GLM parameters were estimated using a robust regression with weighted-least-squares (36).

Contrast images for each of the four conditions compared to the fixation baseline were submitted to the second level 2×2 ANOVA using Gender Type (*Conceptual* and *Formal*) and Gender Congruency (*Match* and *Mismatch*) as factors. This analysis allowed us to check for main effects and interactions. Population-level inferences were tested using a threshold of p < 0.001 uncorrected with a voxel extent higher than 100 voxels such that only those peaks or clusters with a p-value corrected for multiple comparisons using family-wise error rate (FWE; 37) and/or false discovery rate (FDR; 38) were considered to be significant (i.e., p < 0.05 for both). All local maxima are reported in the results tables as MNI coordinates (Evans et al., 1993).

In addition to the classical whole-brain analysis, we also estimated the time course of the hemodynamic response function (HRF) of the spatially unsmoothed time series in a group of Regions of Interest (ROI). The HRF was estimated following the Finite Impulse Response [FIR] (39) implemented in the Marsbar toolbox (http://marsbar.sourceforge.net/download.html), which was then characterized by its maximum amplitude peak of amplitude and the latency corresponding to the location of this maximum. This approach allowed us to determine whether the temporal characterization of the brain response for each ROI complemented in some way the information extracted from the direct comparison between conditions. These analyses also increased the statistical power of each comparison reducing the dimensionality of the problem. The ROIs used were built in MNI space combining functional and anatomical criteria such that all voxels: a) were included in the group-level effect of the contrast *All Condition vs Null;* b) were connected to a local t maxima; and c) were included in one AAL structural ROI.

### Functional connectivity analysis

In the current study, we explored whether the pre-defined ROIs show differential coupling with other brain regions depending on the critical experimental manipulations using generalized psychophysiological interactions (gPPI toolbox; http://www.nitrc.org/projects/gppi, 40). Here a multiregional approach previously used by Cocchi, Zalesky, Fornito and Mattingley (41) was performed (42). This approach included 24 spherical seed regions that were built in MNI space. Each ROI was defined for each participant as the first eigenvariate of the time series of all active voxels within six mm radius spheres centered on the maximum peak of activation resulting from the group-level effect of the F-test contrast *All conditions vs. Null* (p<0.05 FWER corrected at the peak level). Given the 24 ROIs included in these analyses, 276 possible connections per subject and condition were generated.

The gPPI approach allowed us to explore whether the response pattern of a seed region would predict the response pattern in another region dependent on the context, without making any assumptions regarding the relationships between experimental conditions (43). The design matrix used to estimate possible psychophysiological interactions spanned the whole experimental space described above (critical conditions: *Formal Gender Mismatch, Formal Gender Match, Conceptual Gender Mismatch*, and *Conceptual Gender Match*). For each experimental condition, a regressor obtained by convolving the canonical hemodynamic response function with delta functions at stimulus onsets was included as a dependent variable. Similarly, for each experimental condition, the gPPI terms that resulted from convolution of the canonical HRF with the multiplication of the deconvolved neural response of the seed ROI and the task-related time course per condition were also included as explanatory variables. These models also comprised the deconvolved brain response for the corresponding seed ROI and also the six motion-correction parameters as regressors.

After the estimation of each general linear model (per seed ROI, per participant), connectivity matrices were created by averaging the β values of the voxels within each ROI, corresponding to the psychophysiological interaction effects for each condition. In order to identify reliable connections between pairs of regions at the group level, we included the individual connectivity matrices in a second-level analysis, using the network-based statistic toolbox (NBS, 23) following a repeated measures ANOVA design. To make population inferences, the NBS creates a new set of data thresholding (T score > 3) each pair-wise association included in the individual connectivity matrices. The network of connections emerging from this procedure (i.e. exceeding the threshold) was fed into a nonparametric permutation test (10000 random permutations) across participants used to establish the significance of the number of connections of the network assigning a p-value controlled for multiple comparisons (FWER, p < 0.05) (for methodological details see 23, 44 for applications of the NBS toolbox).

For those ROIs resulting as critical to integrate formal and conceptual information (i.e., fMRI interaction effect), individual gPPI contrast images were entered into a second-level 2×2 ANOVA using Gender Type (*Conceptual* and *Formal*) and Gender Congruency (*Match* and *Mismatch*) as factors. Whole-brain correction for multiple comparisons was applied by combining a significance peak-level of p < 0.001 uncorrected, with a voxel extent higher than 100 voxels, such that only those peaks or clusters with a p-value corrected for multiple comparisons using family-wise error rate (FWER; 37) and/or false discovery rate (FDR; 38) were considered to be significant (i.e., p < 0.05 for both).

### Graph-theory properties

The weighted undirected connectivity matrices resulting from the gPPI analysis per pair of regions were further pooled into a matrix for each of the subjects. Nodes and edges were characterized according to six graph-theory properties using the Brain Connectivity Toolbox (brain-connectivity-toolbox.net): 1) node betweenness centrality (i.e., the fraction of all shortest paths that contain a given node); 2) node strength (i.e., the sum of weights of inward and outward links connected to the node); 3) links per node or node degree (i.e., the total number of inward and outward links connected to the node); 4) local efficiency (i.e., the average of inverse shortest path length computed on the neighborhood of the node); 5) weighted clustering coefficient (i.e., the average (geometric mean) of all triangles associated with each node and is equivalent to the fraction of the node’s neighbors that are neighbors of each other); 6) modularity (i.e., subdivision of the network into non-overlapping groups of nodes using the Newman’s community detection algorithm (45)); and 2) betweeness centrality (46). Combining these indexes makes it possible to identify crucial regions characterized by high communication capacity that facilitates functional integration (i.e., the so-called hubs) (14, 15). Differences across conditions were then assessed using a permutation test that provides exact p-values for any number of subjects and estimation algorithms. The distribution of the maximum statistics t was then used to set significance levels that correct for multiple comparisons (47).

## Acknowledgments

This research was supported by the Basque Government through the BERC 2018-2021 program, by the Agencia Estatal de Investigación through BCBL Severo Ochoa excellence accreditation SEV-2015-0490, the Ramon y Cajal Fellowship RYC-2017-21845, and by the Spanish Ministry of Science and Education through project RTI2018-093547-B-I00. We would like to thank BCBL’s Lab Department, in particular David Carcedo which has been working with us during participant selection and data recording processes. We would like to thank also Magda Altman and Brendan Costello for her useful comments on the manuscript.

## Notes

### Competing Interest Statement

The authors have declared no competing interest.

### Summary of Updates

The results section has been updated incorporating new findings concerning the connectivity analysis with the corresponding changes in the discussion section.

