## Supplementary material for "Linguistic input drives brain network configuration during language comprehension": Figure 1S. Clustering indiviadual variability

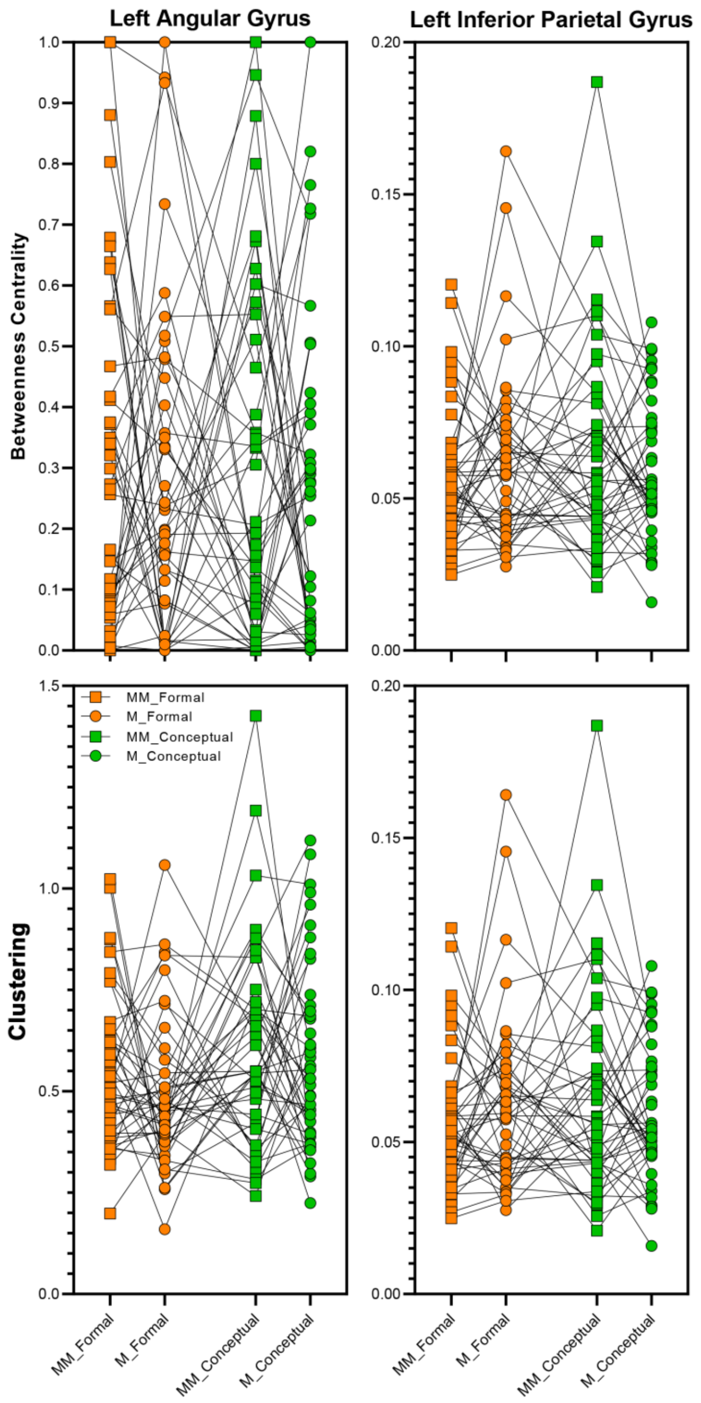


**Figure 1S.** Individual condition-dependent changes in weighted clustering coefficient and betweeness centrality for the angular and inferior parietal gyri. Betweeness centrality (i.e., between 0 and 1) indicates the fraction of all shortest paths that contain a given node. The highest values of betweeness centrality (1) correspond with regions participating in a large number of shortest paths. The weighted clustering coefficient is the average (geometric mean) of all triangles associated with each node and is equivalent to the fraction of the node's neighbors that are neighbors of each other. Notably, there is a huge variability across participants, especially in the betweeness centrality scores. This should be critical for further works.
